# Proteostatic remodeling of small heat shock chaperones - crystallins by Ran-binding protein 2 and the peptidyl-prolyl *cis-trans* isomerase and chaperone activities of its cyclophilin domain

**DOI:** 10.1101/2024.01.26.577462

**Authors:** Hemangi Patil, Kyoung-in Cho, Paulo A. Ferreira

## Abstract

Disturbances in phase transitions and intracellular partitions of nucleocytoplasmic shuttling substrates promote protein aggregation - a hallmark of neurodegenerative diseases. The modular Ran-binding protein 2 (Ranbp2) is a cytosolic molecular hub for rate-limiting steps of disassembly and phase transitions of Ran-GTP-bound protein ensembles exiting nuclear pores. Chaperones also play central roles in phase transitions and proteostasis by suppressing protein aggregation. *Ranbp2* haploinsufficiency promotes the age-dependent neuroprotection of the chorioretina against photo-oxidative stress by proteostatic regulations of Ranbp2 substrates and by countering the build-up of poly-ubiquitylated substrates. Further, the peptidyl-prolyl *cis-trans* isomerase (PPIase) and chaperone activities of the cyclophilin domain (CY) of Ranbp2 modulate the proteostasis of selective neuroprotective substrates, such as hnRNPA2B1, STAT3, HDAC4 or L/M-opsin, while promoting a decline of ubiquitylated substrates. However, links between CY PPIase activity on client substrates and its effect(s) on ubiquitylated substrates are unclear. Here, proteomics of genetically modified mice with deficits of Ranbp2 uncovered the regulation of the small heat shock chaperones – crystallins by Ranbp2 in the chorioretina. Loss of CY PPIase of Ranbp2 up-regulates αA-crystallin proteostasis, which is repressed in non-lenticular tissues. Conversely, the αA-crystallin’s substrates, γ-crystallins, are down-regulated by impairment of CY‘s C-terminal chaperone activity. These CY-dependent effects cause the age-dependent decline of ubiquitylated substrates without overt chorioretinal morphological changes. A model emerges whereby the Ranbp2 CY-dependent remodeling of crystallins’ proteostasis subdues molecular aging and preordains chorioretinal neuroprotection by augmenting the chaperone buffering capacity and the decline of ubiquitylated substrates against proteostatic impairments. Further, CY’s moonlighting activity holds *pan*-therapeutic potential against neurodegeneration.

## INTRODUCTION

The Ran-binding protein 2 (Ranbp2; aka, Nup358) comprises the cytoplasmic filaments emanating from nuclear pores, where it acts as a molecular hub for rate limiting steps of nuclear export [1–6]. Ranbp2 mediates the disassembly of Ran-GTP-bound protein ensembles upon exiting the cytosolic face of nuclear pores. These processes involve phase transitions of Ran-GTP-bound and liberated cargoes [7, 8]. Ranbp2 regulates the nucleocytoplasmic partition and phase transitions of nuclear transport receptors and client substrates elicited by genetic and environmental insults, and by aging [9–13]. Impairments of these biophysical processes are a hallmark of neurodegeneration [1, 14]. These disturbances culminate in the formation of insoluble protein aggregates and membraneless bodies that become resistant or inaccessible to degradation by the ubiquitin-proteasome system (UPS) in neurodegenerative disease [15, 16].

Molecular chaperones have critical but complementary roles in proteostasis under homeostatic and stress environments [17]. Chaperones transiently interact with client substrates to overcome energy barriers in protein biogenesis, to catalyze rate-limiting steps of protein folding or to favor intramolecular interactions that avert the genesis of metastable or kinetically trapped conformers prone to form aggregates of high thermodynamic stability [18–20]. In general, chaperones display significant promiscuity toward client substrates and this feature is postulated to promote cellular fitness by enabling proteostatic adaptations against stressors [21–23]. By contrast, few chaperones have cell-type, tissue or developmental expression or functions that are also accompanied by chaperone activities on selective client substrates. For example, mutations in the ER-resident cyclophilin B (PPIB), which harbors chaperone and peptidyl-prolyl *cis-trans* isomerase (PPIase) activities, causes osteogenesis imperfecta by impairing collagen biogenesis [24–27]. In contrast, mutations in the ER-resident cyclophilin, Nina-A of *Drosophila* specifically affect the biogenesis of a subset of photon-capturing G protein-coupled receptors, opsins, in photoreceptor neurons [28–30]. The molecular bases of the substrate selectivity and evolutionary divergent specificities of these and other cyclophilins remain unresolved [20].

Crystallins belong to a family of small heat shock proteins of conspicuous solubility and stability and apparently, promiscuous chaperone and co-chaperone activities for misfolded proteins [19, 22, 31–36]. In particular, crystallins have exceptional physiochemical properties that enable these chaperones to exceed critical concentration (supersaturation) [37, 38], and negligible turnover and light scattering in the eye lens, where they perform a structural role in providing transparency throughout the human lifespan [39, 40]. For example, γΒ-crystallin concentration in the lens reaches 860 mg/mL without undergoing crystallization and γΒ-crystallin has exceptional stability against harsh chemical and physical agents [41–45]. These unique physiochemical properties of γ-crystallin(s) are possible owing to two other members of small heat shock proteins, αA-crystallin (HSPB4) and αB-crystallin (HSPB5) that chaperone members of the βγ-crystallin family [34, 46–51] and that account for ≈40% of the total protein of the lens [52]. In contrast to monomeric γ-crystallins, α-crystallins form poly-dispersed, dynamic and high-order oligomers that sequester misfolded client substrates [53–58]. α-crystallins are referred to as “holdase” chaperones owing to the capacity of small heat shock proteins to keep unfolded, aggregation-prone and non-native client substrates soluble and in a folding competent state, but without an inherent capacity to refold these *in vitro* [34, 56, 59–61]. Age-related posttranslational modifications and mutations in crystallins promote light-scattering aggregates of crystallins, the opacity of the lens and ultimately, cataracts [62]. Notably, αA-crystallin plays a central role in cataract pathogenesis, because mice with loss of αA-crystallin result in the opacity of the lens, whereas mice with loss of αB-crystallin do not develop cataracts [51, 63, 64]. Although significant insights on the roles of crystallins in lens biology were gained during the past decades, the physiological substrates of crystallins remain scant [36, 49].

αB-crystallin is expressed in lens as well in non-lenticular tissues [65–70]. Although αA-crystallin is primarily expressed in the lens [68, 71, 72], basal levels of αA-crystallin and γ-crystallin are expressed in photoreceptor neurons [73] and across retinal neurons, where they prominently localize to neuronal bodies [74]. Notably, αA-crystallin appears restricted to nuclear membranes of photoreceptors [74]. A role of αA-crystallin in the chorioretina is supported by the observation that human mutations in αA-crystallin or its loss of expression cause micropthalmia [51, 75]. In contrast to αA-crystallin, stressors induce αB-crystallin owing to the presence of a unique heat-shock promoter element in the αB-crystallin gene [76, 77]. The roles and regulation of crystallins and their physiological substrates in non-lenticular tissues, such as the chorioretina, are not understood. Accruing evidence indicates that α-crystallins play roles in neuroprotection, and pathogenesis of neurodegenerative and other diseases, such as diabetes [73, 78–81]. First, crystallins suppress the biogenesis of amorphous and fibrillary deposits, such as amyloid fibrils and α-synuclein [82–86]. Second, heat shock proteins, such as crystallins, are components of drusen deposits in early and advanced stages of age-related macular degeneration (AMD), and they are biomarkers of AMD [87–93]. Finally, crystallins respond to physical insults, such as phototoxicity and deleterious mutations to the chorioretina, where they contribute to anti-apoptotic activity [87, 94–103]. However, the molecular bases of the stress-dependent effects and regulation of crystallins in proteostasis of non-lenticular tissues are obscure.

Our prior research has shown that haploinsufficiency of *Ranbp2* promotes the age-dependent neuroprotection of the chorioretina against phototoxicity by suppressing the damage and apoptosis of photoreceptor neurons and the accumulation of intracellular deposits in the retinal pigment epithelium (RPE) [11, 12]. Among other effects, this neuroprotection is reflected by the suppression of the light-elicited accumulation of poly-ubiquitylated substrates [11]. Subsequent studies found that the loss of PPIase activity of the cyclophilin domain (CY) of Ranbp2 in mice also causes a decrease of poly-ubiquitylated substrates in the retina absent of stressors, while enhancing the visual pathway (to the brain) by light [104]. This is reflected by a decrease of the latency of the implicit times of the light-stimulated visual evoked potential [104]. Further, we found that the CY of Ranbp2 harnesses non-overlapping PPIase and chaperone activities on neuroprotective substrates in mice [104]. Loss of CY PPIase leads to a decrease of heterogeneous nuclear ribonucleoprotein A2B1 (hnRNPA2B1) and to a nuclear shift of histone deacetylase 4 (HDAC4), whereas impairment of the C-terminal chaperone activity of CY enhances the proteostasis of M-opsin [104]. CY also associates to the neuroprotective substrate, signal transducer and activator of transcription 3 (STAT3), and STAT5 [104]. Some of these effects are recapitulated by novel small molecules against the CY PPIase pocket of Ranbp2 [105]. However, a link between the client substrates of Ranbp2 CY and the effects of CY in ubiquitylation of substrates remains unclear.

Here, we report that Ranbp2 also modulates the proteostasis of crystallins and that selective disturbances of PPIase and chaperone activities of CY Ranbp2 promotes the remodeling of crystallins’ proteostasis by elevating the basal levels of αA-crystallin and by decreasing the levels of the client substrates of αA-crystallin, γ-crystallins. This remodeling of crystallins’ proteostasis by loss of the PPIase activity of CY of Ranbp2 is associated with the age-dependent decline of ubiquitylated substrates selectively in the retina and RPE. A model emerges whereby targeting the CY PPIase activity of Ranbp2 preordains the chorioretina against molecular aging and to neuroprotection by enhancing the buffering capacity of crystallins for client substrates. These effects suppress proteotoxicity, while declining the presentation of client substrates to the ubiquitin-proteasome system (UPS) against proteostatic stressors that promote aberrant phase transitions during disassembly of protein ensembles exiting the nuclear pores.

## RESULTS

### Generation of mice with genetic ablation of Ranbp2 and with mutations in modular activities of Ranbp2

In this study, we took advantage of several genetically engineered lines of *Ranbp2* that we have generated and reported in prior studies to identify novel client substrates that are affected by the complete loss of Ranbp2 or modular activities of Ranbp2 [10, 12, 104]. Our ultimate focus was to uncover client substrates affected by loss of the PPIase or C-terminal chaperone activity of CY of Ranbp2 (and herein referred to as moonlighting activity of CY) and that play a central role in proteostasis. We focused on proteostatic impairments affecting the RPE and the retina for several reasons. First, these tissues are vulnerable to environmental insults (e.g. photo-oxidative stress) and predisposed to proteotoxicity and AMD [1]. AMD is a multifactorial disease characterized by the induction of heat shock proteins, the formation of large laminar heterogeneous aggregates (drusen) with aging and by the irreversibly loss of photoreceptors neurons [93, 106, 107]. Second, we have previously shown that haploinsufficiency of *Ranbp2* preordains the neuroprotection of these tissues against proteostatic stressors, such as photo-oxidative stress [11, 12]. Notably, dysregulations of the moonlighting activity of CY of Ranbp2 causes proteostatic imbalances of a restricted set of client substrates with neuroprotection potential and some of these substrates are also affected in mice with insufficiency of Ranbp2 [11, 104]. However, a link between the client substrates affected by the impairment of CY moonlighting activity of Ranbp2, such as STAT3, HDAC4 and hnRNPA2B1, and the observed decline of poly-ubiquitylated substrates caused either by loss of CY moonlighting activity of Ranbp2 in naïve mice or by photo-oxidative stress in *Ranbp2* haploinsufficient mice are unclear [11, 12, 104]. Hence, we postulated that the CY moonlighting activity of Ranbp2 modulates the proteostasis of ubiquitylated substrates by affecting client substrates directly implicated in the regulation of proteostasis or the presentation of substrates to the UPS. This hypothesis is also strengthened by the findings that a domain upstream of CY of Ranbp2, the cyclophilin-like domain (CLD), associates with the subunits of the 19S regulatory cap of the 26S of the proteasome [104, 108–110], and that CLD and CY functions of Ranbp2 may be coupled to regulate proteostasis (figure 1a).

**Figure 1.**
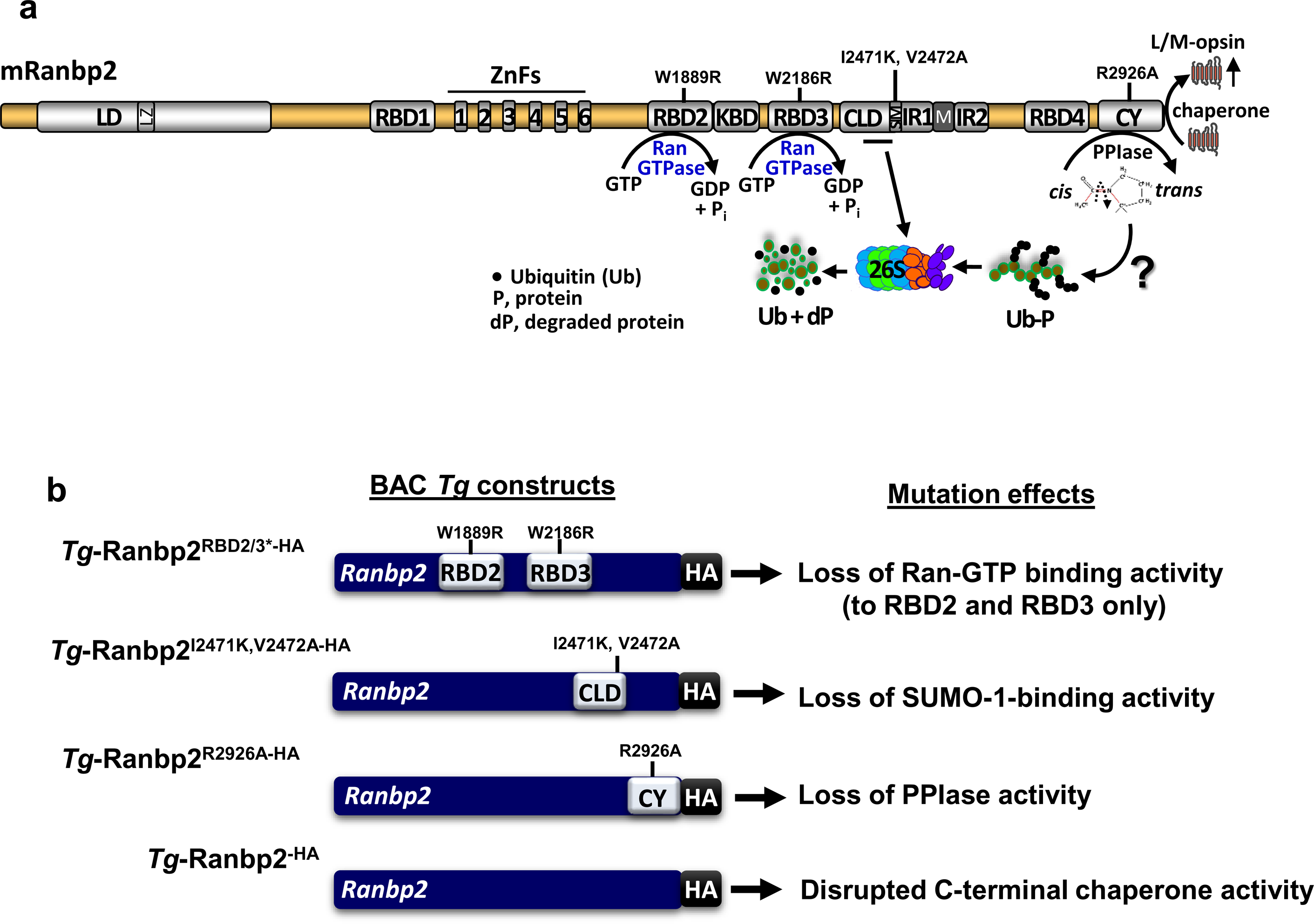
Primary structure and selective modular activities of Ranbp2, and BAC constructs of Ranbp2. (a) Primary structure and domains of murine Ranbp2 with mutations examined by this study in proteostasis. Modular activities of associated to critical residues and mutations of domains of Ranbp2 are shown. RBDs of Ranbp2 bind with high-affinity Ran-GTP and promote its hydrolysis. *Abbreviations*: mRanbp2, murine Ran-binding protein 2; RBD_n1-4_, leucine-rich domain; Ran-binding domain 1-4; LD, ZnFs, zinc-finger-rich domain with ZnF motifs 1 - 6; KBD, kinesin-1-binding domain; CLD, cyclophilin-like domain; IR1and IR2, internal repeats 1 and 2; SIM, SUMO-1-interacting (binding) motif; CY, cyclophilin domain; Ran, RAs-related Nuclear protein; PPIase, peptidyl-prolyl *cis-trans* isomerase; 26S, 26 proteasome; ?, interrogation by this study of the effects of PPIase and C-terminal chaperone activities of CY (and CLD) in proteostasis and ubiquitylated substrates. Relative scales of domains of Ranbp2 are shown. (b) Designation of transgenic constructs of bacterial artificial chromosomes (BAC) containing the complete *Ranbp2* gene with different mutations as depicted in (a). The known biochemical effects of the mutations are shown (right).

In this study, we employed wild type mice and the following six genetically engineered lines of *Ranbp2*: *a*) mice with genetic loss of *Ranbp2* selectively in the RPE (RPE-cre::*Ranbp2^-/-^*; herein referred as RPE-cre::*-/-*) [10]; *b*) mice with the constitutive loss of *Ranbp2* (*Ranbp2^-/-^*;; herein referred as *-/-*) [12, 104, 111]; *c*) mice with a bacterial artificial chromosome (BAC) transgene (*Tg*) of the complete *Ranbp2* gene with the point mutations W1889R and W2186R in the respective Ran-binding domain 2 (RBD2) and RBD3 and that cause the uncoupling of Ran-GTP-binding (*Tg*-Ranbp2^RBD2/3*-HA^; figures 1a and 1b); *d*) mice with a BAC *Tg* of the complete *Ranbp2* gene with the point mutations, I2471K and V2472A, in the C-terminal region of CLD and overlapping internal repeat-1 (IR1) of Ranbp2, and that abolish the SUMO-1-consensus interacting motif (SIM) [104] (*Tg*-Ranbp2^I2471K,V2472A-HA^; figures 1a and 1b); *e*) mice with a BAC *Tg* of the complete *Ranbp2* gene, which harbors the point mutation R2926A causing the catalytic loss of PPIase (*Tg*-Ranbp2^R2926A-HA^; figures 1a and 1b); and *f*) mice with a BAC *Tg* of the complete wild-type *Ranbp2* gene and that contains the insertion of the influenza hemagglutinin (HA)-tag at the C-terminal end of CY of Ranbp2 and that disrupts the C-terminal chaperone activity of CY (*Tg*-Ranbp2^-HA^; figures 1a and 1b). All BAC *Tg* lines with the point mutations described contained also the HA-tag insertion at the C-terminal end of CY of Ranbp2 (figure 1b). Because mice with constitutive loss of *Ranbp2* are embryonic lethal, BAC *Tg* of Ranbp2 with point mutations were placed in a tissue (RPE)-selective or constitutive *null Ranbp2* (*-/-*) genetic background, and as described elsewhere [10, 104]. Collectively, these lines were used to compare and ultimately, to uncover by proteomics targets with known and direct roles in proteostasis and with target impairments retained by *Tg*-Ranbp2^R2926A-HA^::-/-, *Tg*-Ranbp2^-HA^::-/- or both.

### Identification of small heat shock proteins - crystallins, differentially expressed between wild-type, RPE-cre::-/- and Tg-Ranbp2^RBD2/3*-HA^::RPE-cre::-/-

In this study, we employed a proteomic approach with two-dimensional difference gel electrophoresis (2D DIGE) to resolve, profile and compare the proteomes of wild type and genetically modified mice of *Ranbp2*. To this effect, tissue lysates were labeled with the CyDye-fluorescent dyes, Cy2, Cy3 and Cy5 in a genotype-dependent manner. This approach was used to first identify co-resolved proteins (spots) with similar pI, apparent molecular weight and electrophoretic mobilities between genotypes. We began by comparing spots between wild type mice, mice with the loss of *Ranbp2* expression selectively in the RPE and mice with the expression of *Tg*-*Ranbp2^RBD2/3*-HA^* in a *null Ranbp2* background in the RPE. The reason for including the *Tg-Ranbp2^RBD2/3*-HA^::RPE-cre::-/-* in the initial analysis is because this genotype recapitulates with a high-degree of fidelity the pathophysiological phenotypes of *RPE-cre::-/-* and it promotes the up-regulation of some components of the Ranbp2 ensemble or client substrates [10]. Protein identities of selected spots were carried out by mass spectrometry (MALDI-TOF/TOF) of mass spectra of tryptic peptides and peptide fragmentation mapping of trypsin-digested spots [10, 104]. We first used RPE of mice at post-natal day 14 (P14), because this developmental stage marks the end of cell division of developing RPE [10, 112], the opening of the eyelids, and the beginning of regional morphological changes in the RPE of both *RPE-cre::-/-* and Tg-*Ranbp2^RBD2/3*-HA^::RPE-cre::-/-* [10].

The proteomic approaches employed with wild-type*, RPE-cre::RPE-cre::-/-* and *Tg-Ranbp2^RBD2/3*-HA^::RPE-cre::-/-* lines, and with other lines described later in this study, led to the consistent identification of four to five co-resolving spots in homogenates of RPE at P14. These comprised spot 1, spot 2, spot 3, clustered spots 4A and 4B, and spot 5 (figure 2a). Among these spots and when compared to wild type mice, the intensity of spots 4A, 4B, and 5 were increased in Tg-*Ranbp2^RBD2/3*-HA^::RPE-cre::-/-* and decreased in *RPE-cre::RPE-cre::-/-,* whereas spots 1 and 2 appeared unchanged between genotypes (figure 2a). Searches of NCBI and SwissProt databases of combined MS and MS/MS spectra with the MASCOT search engine generated protein and total ion scores with confidence intervals (C.I.) of 100% for the following proteins (figure 2b). Spots 1 and 2 with similar pI but an apparent difference in molecular mass of 2-3 kDa comprised αA-crystallin (figure 2b). The αA-crystallin isoform of higher molecular weight is likely generated from of a splice variant of αA-crystallin, which contains an insert encoding 23-amino acids in *Mus musculus* [74, 113, 114]. By contrast, spots 3, 4A, 4B and 5 comprised several highly homologous members of γ-crystallin family of proteins [115]. Each spot had two or more members of γ-crystallin with very closely related molecular masses and pI (figure 2b). The exception was spot 5, which was represented only by γS-crystallin with an observed pI of ≈6.8, which was distinct from the other γ-crystallin members. Despite slight variations in molecular masses between spots 3, 4A, 4 B, each of these spots shared two members of γ-crystallins, such as γA-crystallin and γC-crystallin. These small electrophoretic mobility variations may reflect limited proteolytic processing, post-translation modification(s) or both of some members of γ-crystallins.

**Figure 2.**
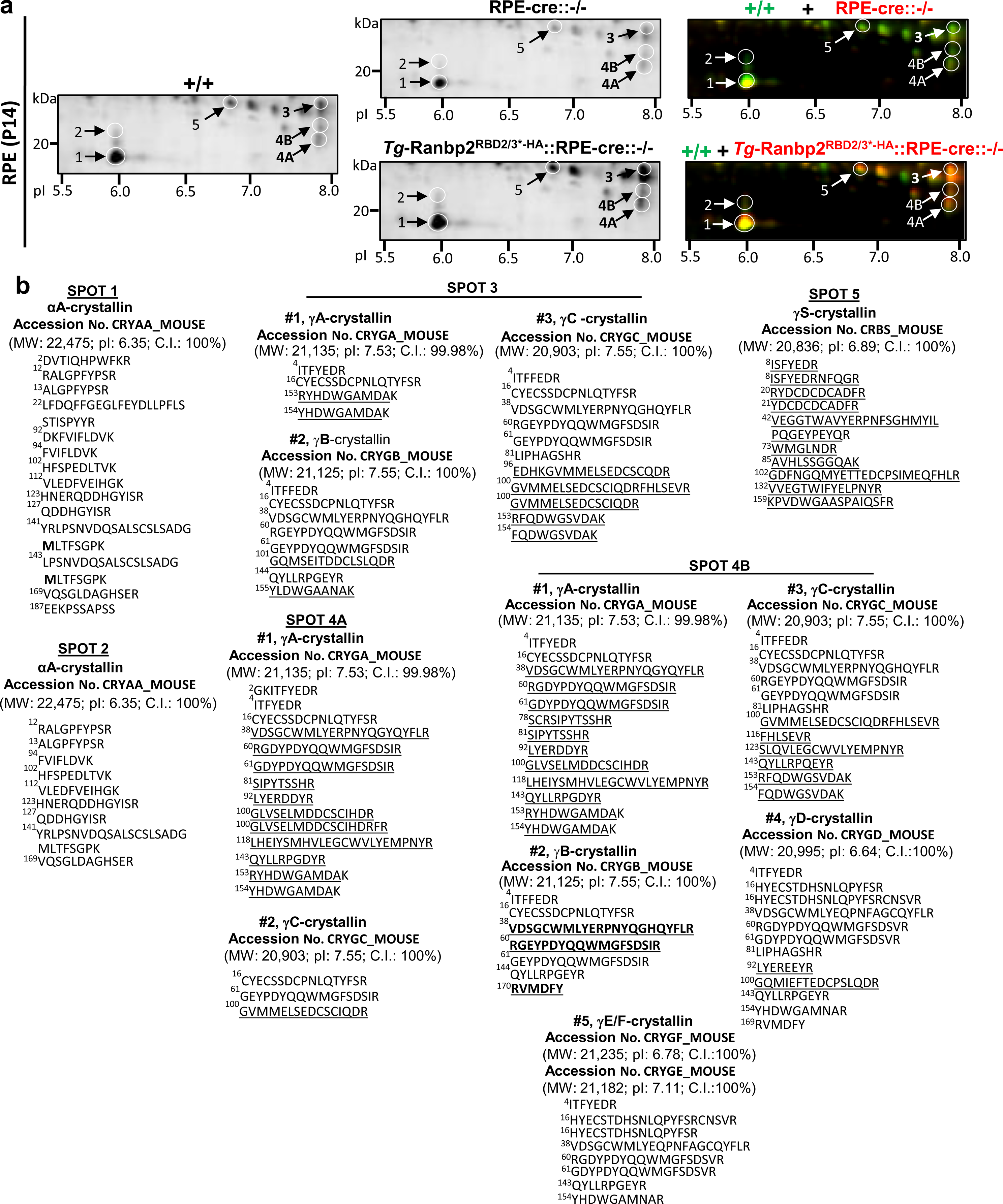
2D-DIGE and tryptic peptides identified by mass spectrometry of RPE of wild type, RPE-cre::-/- and *Tg*-Ranbp2^RBD2/3*-HA^::RPE-cre::-/- mice at P14 of age. (a) Proteomic differences between RPE of age-matched wild type (+/+), RPE-cre::-/- and *Tg*-Ranbp2^RBD2/3*-HA^::RPE-cre::-/- mice resolved by isoelectric focusing and SDS-PAGE. Homogenates of wild type (+/+), RPE- cre::-/- and *Tg*-Ranbp2^RBD2/3*-HA^::RPE-cre::-/- were labeled with cyanine dyes, Cy2, Cy3 and Cy5. A set of selected spots of defined apparent molecular masses and isoelectric focusing points (pI) were marked (circles) and identified by mass spectrometry (as shown below in (b)). Right panels are overlays of 2D-DIGE between two genotypes. (b) Identification of spots marked in (a) from mass spectra of tryptic peptides and peptide fragmentation mapping by MALDI-TOF/TOF and Mascot searches. Search results with a confidence of interval (C.I.) of 100% of protein and total ion scores from MS and MS/MS spectra are shown. GenBank accession numbers, predicted molecular weights and predicted pI of crystallins identified are shown. Sequences unique to a γ-crystallin isoform are underlined. Because of the high homology between members of γ-crystallin family, sequences shown in bold and underlined were combined sometimes to assign a crystallin isotype. *Abbreviations*: RPE-cre::*-/-,* RPE-cre::*Ranbp2^-/-^* (mice with the endogenous loss of *Ranbp2* selectively in the RPE); *Tg*-Ranbp2^RBD2/3*-HA^::RPE-cre::*-/-*, mice expressing *Tg*-Ranbp2^RBD2/3*-HA^ (described figure 1b) in RPE without endogenous *Ranbp2*; RPE, retinal pigment epithelium; 2D DIGE, two-dimensional difference gel electrophoresis.

### Immunochemical quantification of the effects of RPE-cre::RPE-cre::-/- and Tg-Ranbp2^RBD2/3*-HA^::RPE-cre::-/- in the proteostasis of crystallins isoforms

We then employed an immunoassay approach to validate independently the identities of the crystallin isoforms and to quantify the relative levels of the crystallin isoforms identified by proteomics in the RPE between genotypes. Homogenates of RPE were resolved by SDS-PAGE and immunoblots were probed with antibodies against αA-crystallin, γ-crystallins, and βΒ2-crystallin. The relative levels of crystallins between genotypes were quantified by densitometric analysis on the respective immunobands of the expected molecular weight. As shown in figure 3a and 3b, αA-crystallin level was unchanged between any genotype. By contrast, and compared to wild type mice, the levels of γ-crystallins, which run as a close doublet, were significantly and strongly increased by 10-fold in *Ranbp2^RBD2/3*-HA^::RPE-cre::-/-* and decreased by over 5-fold in *RPE-cre::RPE-cre::-/-*. We also examined the levels of βΒ-crystallin, which is a member of the βγ-crystallin family of proteins. The levels of βΒ2-crystallin directionally and significantly recapitulated the directional changes observed for γ-crystallins between genotypes. Hence, these observations support a role of Ranbp2 and selective modular activities of Ranbp2 in the proteostatic regulation of a subset of members of the crystallin family of proteins in the RPE at P14. In particular, loss of Ranbp2 selectively suppresses the proteostasis of γ-crystallins and βΒ2-crystallin, whereas loss of the Ran-GTP-binding activity of RBD2 and RBD3 of Ranbp2 has the opposite effect by promoting the strong increase of these crystallins in the RPE at P14.

**Figure 3.**
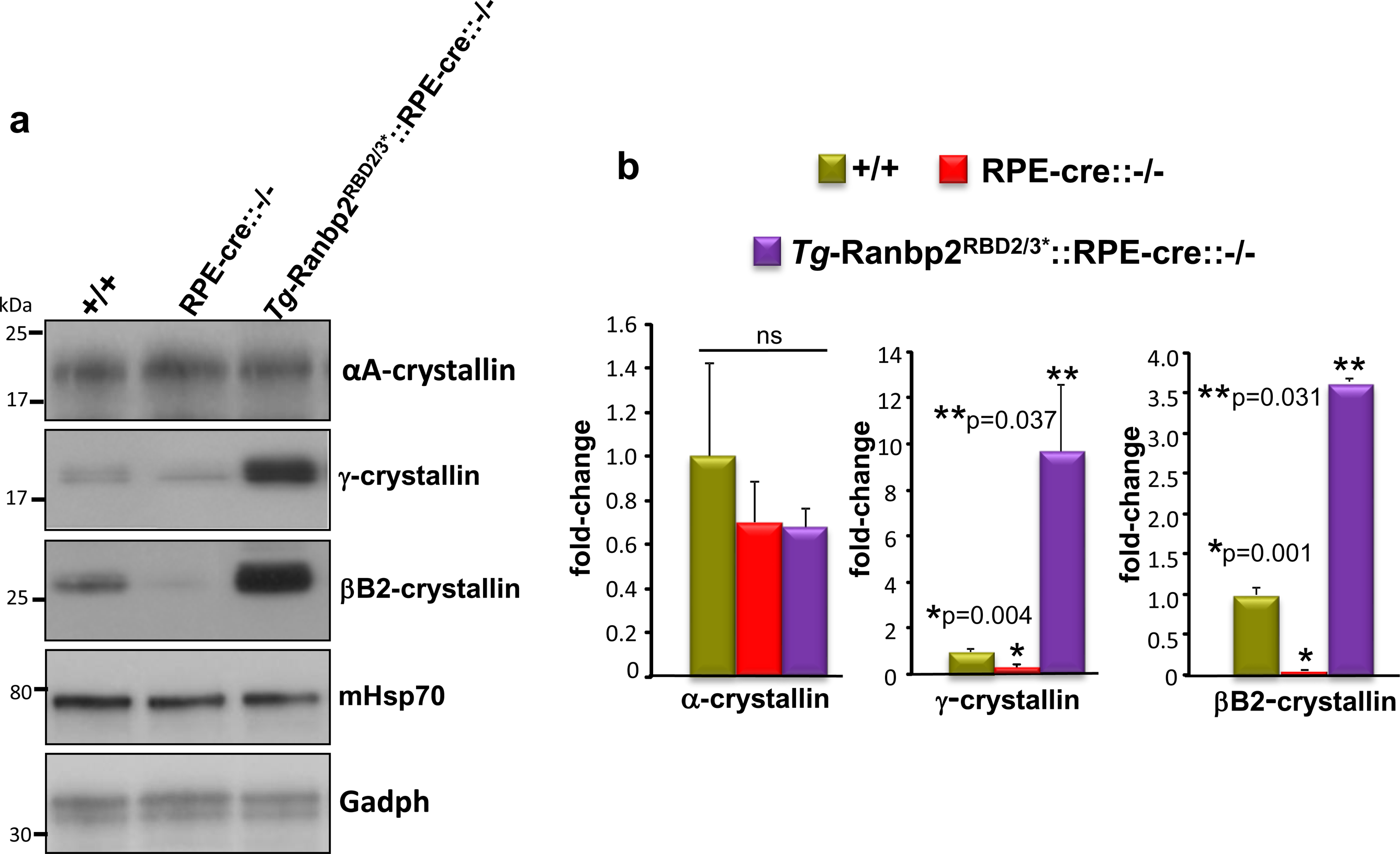
Relative expression levels of crystallins isotypes in RPE homogenates of wild type, RPE-cre::-/- and *Tg*-Ranbp2^RBD2/3*-HA^ mice at P14 of age. (a) Immunoblot of RPE homogenates of age-matched wild type (+/+), RPE-cre::-/- and *Tg*-Ranbp2^RBD2/3*-HA^::RPE-cre::*-/-* probed with antibodies against αA-crystallin, βB2-crystallin and γ-crystallins. mHsp70 and Gadph are loading and housekeeping protein controls. γ-crystallins resolved as a doublet. (b) Relative levels of αA-crystallin, βB2-crystallin and γ-crystallins between RPE of wild type (+/+), RPE-cre::-/- and *Tg*-Ranbp2^RBD2/3*-HA^::RPE-cre::*-/-* mice at P14 of age. Data represent the mean ± S.D (error bars), *n*=4 mice. *Abbreviations*: RPE, retinal pigment epithelium.

### Age-dependent suppression of crystallins in mature RPE and up-regulation of crystallins in Tg-Ranbp2^R2926A-HA^::-/- and Tg-Ranbp2^I2471K,V2472A-HA^::-/-

As noted earlier, crystallins are regulated developmentally and the CLD and the moonlighting activity of CY of Ranbp2 modulate protein homeostasis (figure 1a). Our goal is to understand the role of the moonlighting activity of CY of Ranbp2 in the mature chorioretina (e.g., retina and RPE). Hence, we employed the same proteomic approaches described earlier by examining and by comparing the proteostasis of αA-crystallin and γ-crystallins approximately to the corresponding spots 1, 2, 3 and 4, since γ-crystallins are reported as substrates for αA-crystallin. We began by examining the RPE of 5-week old mice between wild type, *Tg*-*Ranbp2^R2926A-HA^::-/- and Tg-Ranbp2^I2471K,V2472A-HA^::-/-*. As we previously reported, naive mice of these genotypes and age do not present overt morphological phenotypes of the chorioretina [104, 116]. As shown in figure 4a, the signal of spots 1, 3 and 4 were strongly decreased in homogenates of RPE of 5-week old wild type mice, and spot 2 was not detected compared to RPE of P14 wild type mice (figure 2a). By contrast and compared to wild type mice, spots 1 and 3 were strongly increased in both *Tg*-*Ranbp2^R2926A-HA^::-/- and Tg-Ranbp2^I2471K,V2472A-HA^::-/-*, whereas spot 2 was modestly increased (figure 4a).

**Figure 4.**
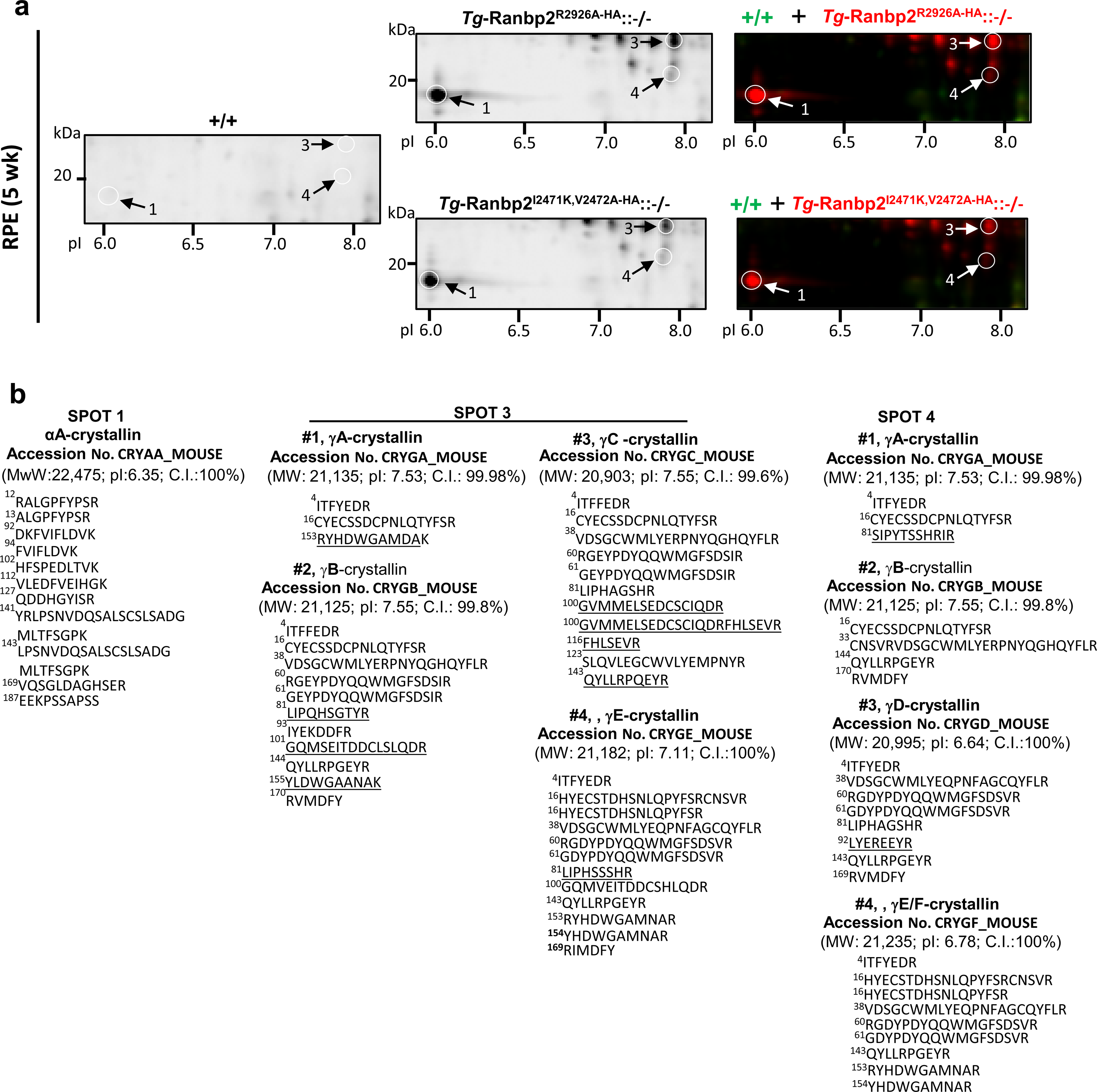
2D-DIGE and mass spectrometry of RPE lysates of wild type, *Tg*-*Ranbp2^R2926A-HA^::-/- and Tg-Ranbp2^I2471K,V2472A-HA^::-/-* at 5-weeks of age. (a) Proteomic differences between RPE of age-matched wild type (+/+), *Tg*-*Ranbp2^R2926A-HA^::-/- and Tg-Ranbp2^I2471K,V2472A-HA^::-/-* mice resolved by isoelectric focusing and SDS-PAGE. RPE homogenates of wild type (+/+), *Tg*-*Ranbp2^R2926A-HA^::-/- and Tg-Ranbp2^I2471K,V2472A-HA^::-/-* were labeled with cyanine dyes, Cy2, Cy3 and Cy5. A set of selected spots of equivalent electrophoretic and isoelectric focusing (pI) migration of figure 2a were marked (circles) and identified by mass spectrometry (as shown below in (b)). Right panels are overlays of 2D-DIGE between two genotypes. (b) Identification of spots marked in (a) from mass spectra of tryptic peptides and peptide fragmentation mapping by MALDI-TOF/TOF and Mascot searches. Search results with a confidence of interval (C.I.) of 100% of protein and total ion scores from MS and MS/MS spectra are shown. GenBank accession numbers, predicted molecular weights and predicted pI of crystallins identified are shown. Sequences unique to a γ-crystallin isoform are underlined. *Abbreviations*: *-/-, Ranbp2^-/-^*; *Tg*-*Ranbp2^R2926A-HA^::-/-*, mice expressing *Tg*-*Ranbp2^R2926A-HA^*(described figure 1b) in a genetic background with the endogenous and constitutive loss of *Ranbp2*; *Tg-Ranbp2^I2471K,V2472A-HA^::-/-*, mice expressing *Tg-Ranbp2^I2471K,V2472A-HA^* (described figure 1b) in a genetic background with the endogenous and constitutive loss of *Ranbp2*; RPE, retinal pigment epithelium; 2D DIGE, two-dimensional difference gel electrophoresis.

We then established the identity of the protein species of spots 1, 3 and 4 by MS/MS and by searches of NCBI and SwissProt databases as described earlier. Protein and total ion scores with confidence intervals (C.I.) of 100% from proteomics of spots 1, 3 and 4 confirmed that spot 1 comprised αA-crystallin, whereas spots 3 and 4 comprised different members of the γ-crystallin family (figure 4b). Spot 3 of RPE of 5-week old wild type mice contained the same γ-crystallin members - γA-crystallin, γB-crystallin and γC-crystallin, as spot 3 of P14 RPE, and another γ-crystallin member, γD-crystallin (figure 4b). By contrast, γC-crystallin was the only crystallin identified in spot 4 of 5-week old wild type mice (figure 4b). We then extended the same 2D-DIGE and proteomic approach and analysis to equivalent spots identified in retinas of age-matched mice. Like in age-matched RPE, spots 1, 2, 3 and 4 were also barely visible in homogenates of retina of 5-week old wild type mice (figure 5a). By contrast, spots 1 and 3, and to a less extent, spots 2 and 4 were strongly increased in homogenates of retinae of *Tg*-*Ranbp2^R2926A-HA^::-/-* and *Tg-Ranbp2^I2471K,V2472A-HA^::-/-* mice (figure 5a). MS/MS found that spots 1 and 2 consisted of αA-crystallin alone, whereas spots 3 and 4 comprised members of the γ-crystallin family. Specifically, spot 3 comprised four members of γ-crystallin - γA-crystallin, γB-crystallin, γC-crystallin and γD-crystallin, whereas spot 4 contained only γC-crystallin (figure 5b). Hence, the RPE and retina of 5-week old wild type mice express basal levels of αA-crystallin and members of γ-crystallin and these crystallins are up-regulated in *Tg*-*Ranbp2^R2926A-HA^::-/-* and *Tg-Ranbp2^I2471K,V2472A-HA^::-/-* mice.

**Figure 5.**
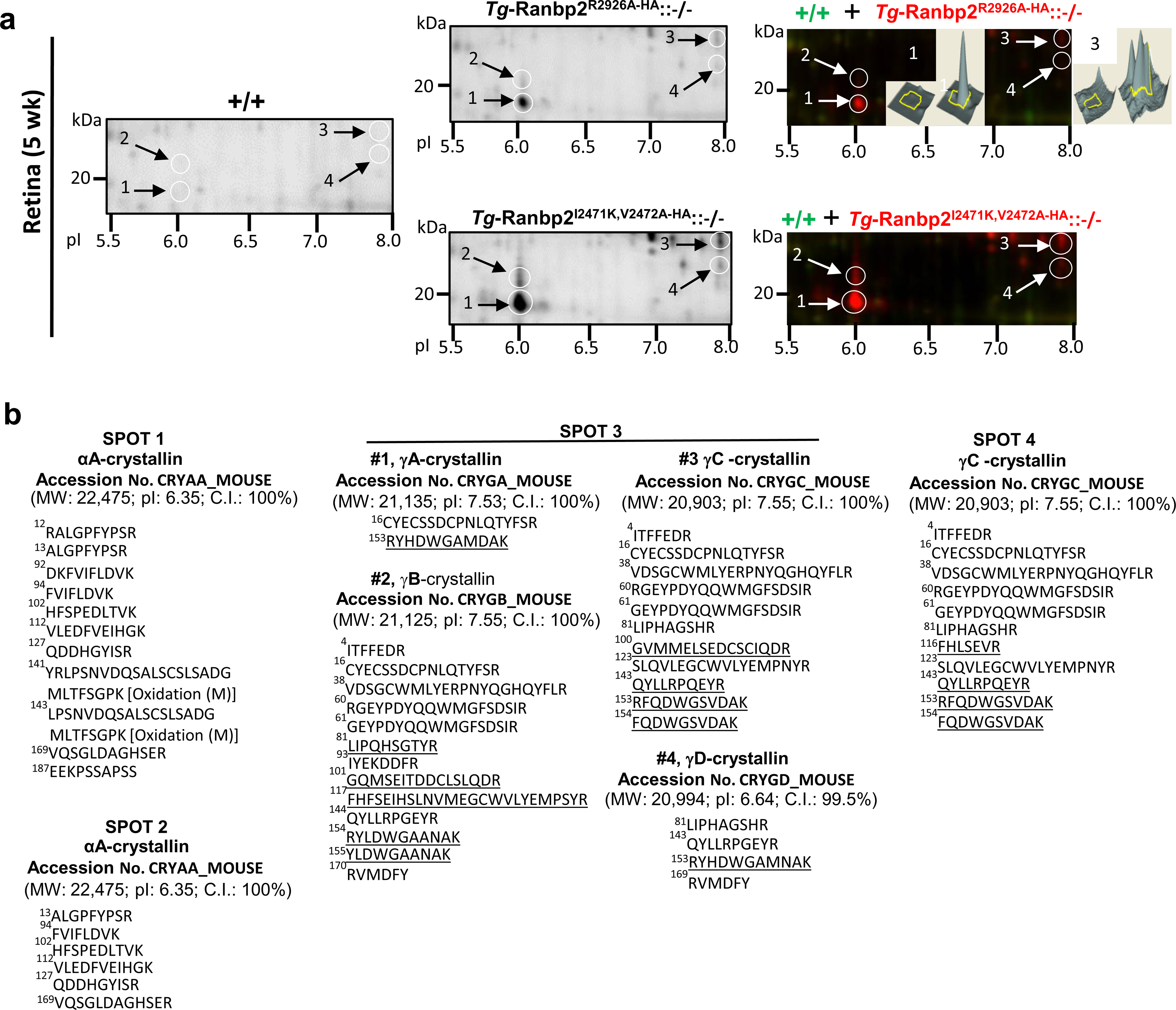
2D-DIGE and mass spectrometry of retinae of wild type, *Tg*-*Ranbp2^R2926A-HA^::-/- and Tg-Ranbp2^I2471K,V2472A-HA^::-/-* at 5-weeks of age. (a) Proteomic differences between retinae of age-matched wild type (+/+), *Tg*-*Ranbp2^R2926A-HA^::-/- and Tg-Ranbp2^I2471K,V2472A-HA^::-/-* mice resolved by isoelectric focusing and SDS-PAGE. Retinal homogenates of wild type (+/+), *Tg*-*Ranbp2^R2926A-HA^::-/- and Tg-Ranbp2^I2471K,V2472A-HA^::-/-* were labeled with cyanine dyes, Cy2, Cy3 and Cy5. A set of selected spots of equivalent electrophoretic and isoelectric focusing (pI) migration of figure 4a were marked (circles) and identified by mass spectrometry (as shown below in (b)). Right panels are overlays of 2D-DIGE between two genotypes. (b) Identification of spots marked in (a) from mass spectra of tryptic peptides and peptide fragmentation mapping by MALDI-TOF/TOF and Mascot searches. Search results with a confidence of interval (C.I.) of 100% of protein and total ion scores from MS and MS/MS spectra are shown. GenBank accession numbers, predicted molecular weights and predicted pI of crystallins identified are shown. Sequences unique to a γ-crystallin isoform are underlined. *Abbreviations*: Genotypes of mice were described in figure 4.

### Immunochemical quantification of loss of PPIase and C-terminal chaperone activities of CY of Ranbp2 in the physiological proteostasis of αA-crystallins and γ-crystallins

As noted earlier, all mouse lines contain an HA-tag at the C-terminal end of CY of Ranbp2. We have previously shown that CY of Ranbp2 has non-overlapping PPIase and C-terminal chaperone activities [104]. To date, only one substrate, L/M-opsin, is known to be affected by a disturbance in the C-terminal chaperone activity of CY, whereas three substrates, hnRNPA2B1, HDAC4 and the UPS activity, are known to be affected by the loss of PPIase activity of CY [104]. Hence, we used quantitative immunochemical assays to examine and compare the effects on the proteostasis of αA-crystallins and γ-crystallins on the following lines: i) *Tg*-*Ranbp2^R2926A-HA^::-/-* mice, which harbor both the loss of PPIase and C-terminal chaperone activities of CY of Ranbp2; ii) *Tg*-*Ranbp2^-HA^::-/-*mice, which harbor only a disturbance of the C-terminal chaperone activity of CY; and iii) wild-type mice with no impairments in CY moonlighting activity (figure 1B).

As shown in figures 6a and 6b, *Tg*-*Ranbp2^R2926A-HA^::-/-* mice presented a significant increase of two αA-crystallin isoforms of apparent molecular weights of ≈20 kDa (≈2.2-fold) and ≈23 kDa (≈6-fold) compared to *Tg*-*Ranbp2^-HA^::-/-* and wild type mice, whereas no significant differences of the level of these αA-crystallin isoforms were observed between *Tg*-*Ranbp2^-HA^::-/-* and wild type mice. The slower electrophoretic mobility isoform, αA-crystallin^SpV^, is reported be a product of spliced variant of αA-crystallin expressed in the mouse retina [74, 113, 114]. As it relates to γ-crystallin proteostasis, the expression of two γ-crystallin isoforms of apparent molecular weights of ≈21 kDa and ≈23 kDa were observed in all mouse lines (figure 6). Notably, there was no difference in the level of the γ-crystallin isoform with the lower molecular weight, γ-crystallin^L^, between *Tg*-*Ranbp2^R2926A-HA^::-/-*, *Tg*-*Ranbp2^-HA^::-/-* and wild type mice. In contrast, and in comparison to wild type mice, the γ-crystallin isoform of higher molecular weight, γ-crystallin^H^, was significantly decreased in both *Tg*-*Ranbp2^R2926A-HA^::-/-* (≈0.4-fold) and *Tg*-*Ranbp2^-HA^::-/-* (≈0.3-fold). Neither the levels of cytosolic heat shock protein 70 (hsc70) and Gadph, nor the levels of nuclear shuttling substrates, COUP-TFI and Nr2e3, partaking in the retinal Ranbp2 ensemble [11] were significantly affected between any mouse line (figure 6). Hence, the physiological and selective loss of PPIase activity of CY of Ranbp2 is required to promote an increase of αA-crystallin isoforms, whereas a disturbance of the C-terminal chaperone activity of CY of Ranbp2 suffices to promote a selective decrease of the γ-crystallin^H^ isoform in the retina.

**Figure 6.**
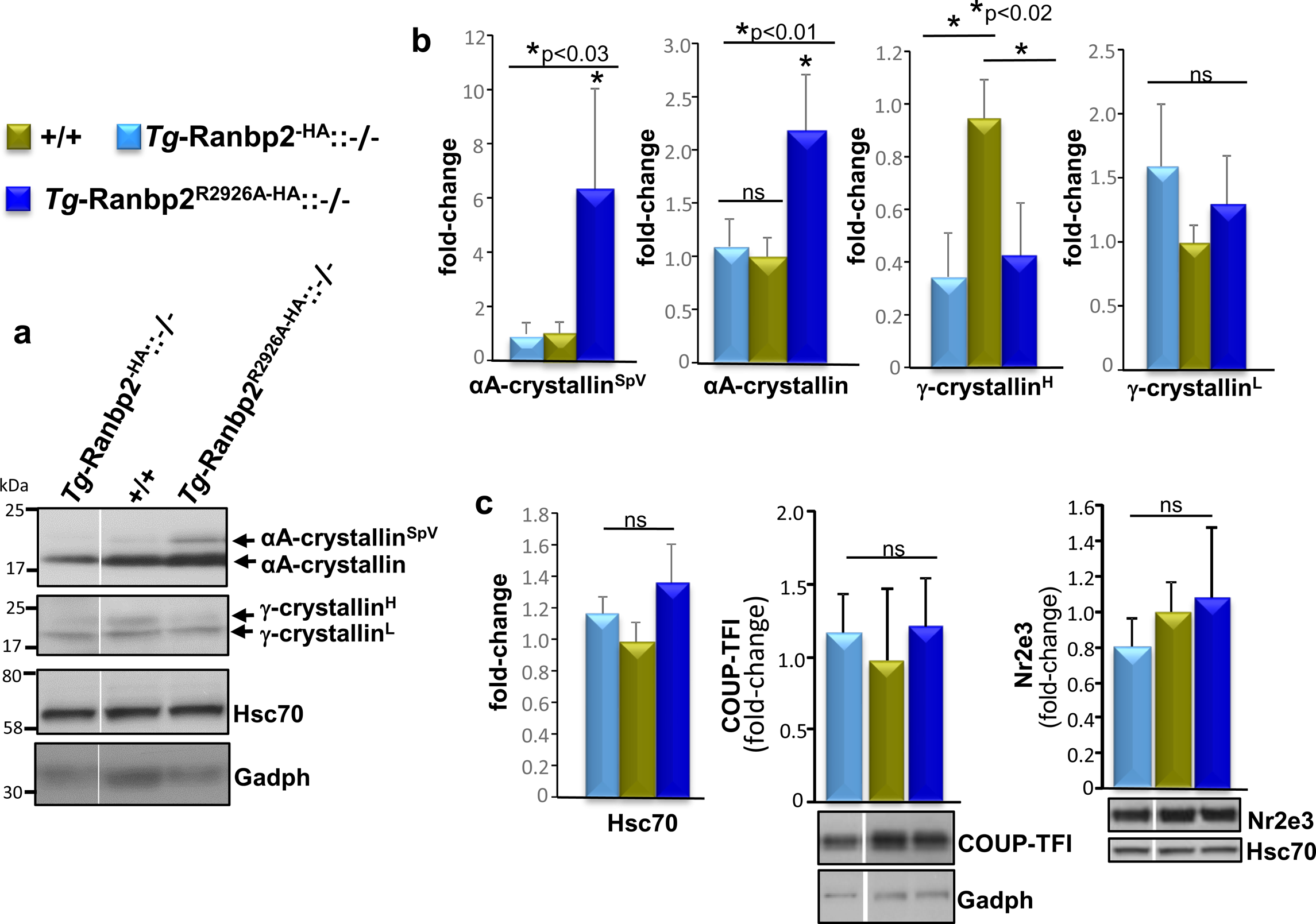
Relative levels of αA-crystallins and γ-crystallins in retinae of wild type, *Tg*-*Ranbp2^R2926A-HA^::-/-* and *Tg-Ranbp2^-HA^::-/-* mice at 5-weeks of age. (a) Immunoblot of retinal homogenates of age-matched wild type (+/+), *Tg*-*Ranbp2^R2926A-HA^::-/- and Tg-Ranbp2^-HA^::-/-* mice probed with antibodies against αA-crystallin and γ-crystallins. Hsc70 and Gadph are loading and housekeeping protein controls. γ-crystallins resolved as a doublet, herein designated γ-crystallin^H^ and γ-crystallin^L^. (b) Relative levels of αA-crystallin and γ-crystallin isoforms between retinal homogenates of wild type (+/+), *Tg*-*Ranbp2^R2926A-HA^::-/- and Tg-Ranbp2^-HA^::-/-* mice at 5-weeks of age. Data represent the mean ± S.D (error bars), *n*=4 mice. *Abbreviations*: SpV, splice variant product; γ-crystallin^H^ and γ-crystallin^L^ are γ-crystallin isoforms of apparent and relative high and low molecular weights. (c) Relative expression levels of Hsc70, COUP-TFI and Nr2e3 between retinal homogenates of wild type (+/+), *Tg*-*Ranbp2^R2926A-HA^::-/- and Tg-Ranbp2^-HA^::-/-* mice at 5-weeks of age. Hsc70 and Gadph are loading and housekeeping protein controls. Data represent the mean ± S.D (error bars), *n*=4 mice. *Abbreviations*: n.s., not significant.

### Age-dependent and selective decrease of ubiquitylated substrates in retina and RPE of Tg-Ranbp2^R2926A-HA^::-/- mice

We have previously shown that mice with a loss of PPIase and a disturbance of C-terminal chaperone activity of CY of Ranbp2 present a decrease of ubiquitylated substrates of a wide range of molecular weights in chorioretinal tissues [104]. The current study indicates that the increase of the proteostasis of αA-crystallin isoforms in *Tg*-*Ranbp2^R2926A-HA^::-/-* likely promotes a proteostatic environment with higher buffering capacity for misfolded or impaired proteins in the chorioretina. It is unclear whether *Tg-Ranbp2^I2471K,V2472A-HA^::-/-* mice, which also present high levels of αA-crystallins and γ-crystallins in the RPE and retina, will also promote proteostasis, even though prior studies indicate that the levels of ubiquitylated substrates are unaffected between wild type and *Tg-Ranbp2^I2471K,V2472A-HA^::-/-* mice of young age [104]. Changes in the levels of ubiquitin and ubiquitylated substrates are a proxy for the activity of the UPS as well as of overall proteostasis owing to changes in the degradation of misfolded and damaged proteins covalently modified by ubiquitin as a result from changes in chaperone activities. Hence, we used an independent and quantitative immunoassay to measure the levels of ubiquitylated substrates in retinae and RPE and with aging between wild type, *Tg*-*Ranbp2^R2926A-HA^::-/-* and *Tg-Ranbp2^I2471K,V2472A-HA^::-/-* mice. We also carried out similar assays with liver tissue of these mice, because of the very low level of expression of αA-crystallin in this tissue [117] and of a clear lack of association of CLD of Ranbp2 with subunits of the 19S cap of the 26S proteasome in liver extracts compared to retinal extracts [108, 109].

As shown in figure 7, there were no significant differences in the concentration of ubiquitylated substrates in the retina and RPE between age-matched wild type and *Tg*-*Ranbp2^R2926A-HA^::-/-* at 5-weeks of age (figure 7a). At 24 and 36-week of age however, the levels of ubiquitylated substrates in the retina and RPE of *Tg*-*Ranbp2^R2926A-HA^::-/-* were lower than wild type mice (figure 7a). Further, the decline of the levels of ubiquitylated substrates became significantly more accentuated with aging in *Tg*-*Ranbp2^R2926A-HA^::-/-*, whereas no significant differences were found in the retina and RPE of wild type mice with aging (figure 7a). Notably, the levels of ubiquitylated substrates did not change in the liver and with aging between genotypes (figure 7a). Parallel examinations of the levels of ubiquitylated substrates in the retina and RPE of wild type and *Tg-Ranbp2^I2471K,V2472A-HA^::-/-* mice also showed no significant differences between genotypes with aging (figure 7b). Hence, neither the loss of SUMO-1-binding function of CLD, nor a disturbance of the C-terminal chaperone activity of CY of Ranbp2 play a role in the decline of ubiquitylated substrates with aging in contrast to *Tg*-*Ranbp2^R2926A-HA^::-/-* (figures 7a and 7b). Further, the loss of PPIase activity in *Tg*-*Ranbp2^R2926A-HA^::-/-* exerts a tissue-selective effect in the retina and RPE, but not in the liver with aging.

**Figure 7.**
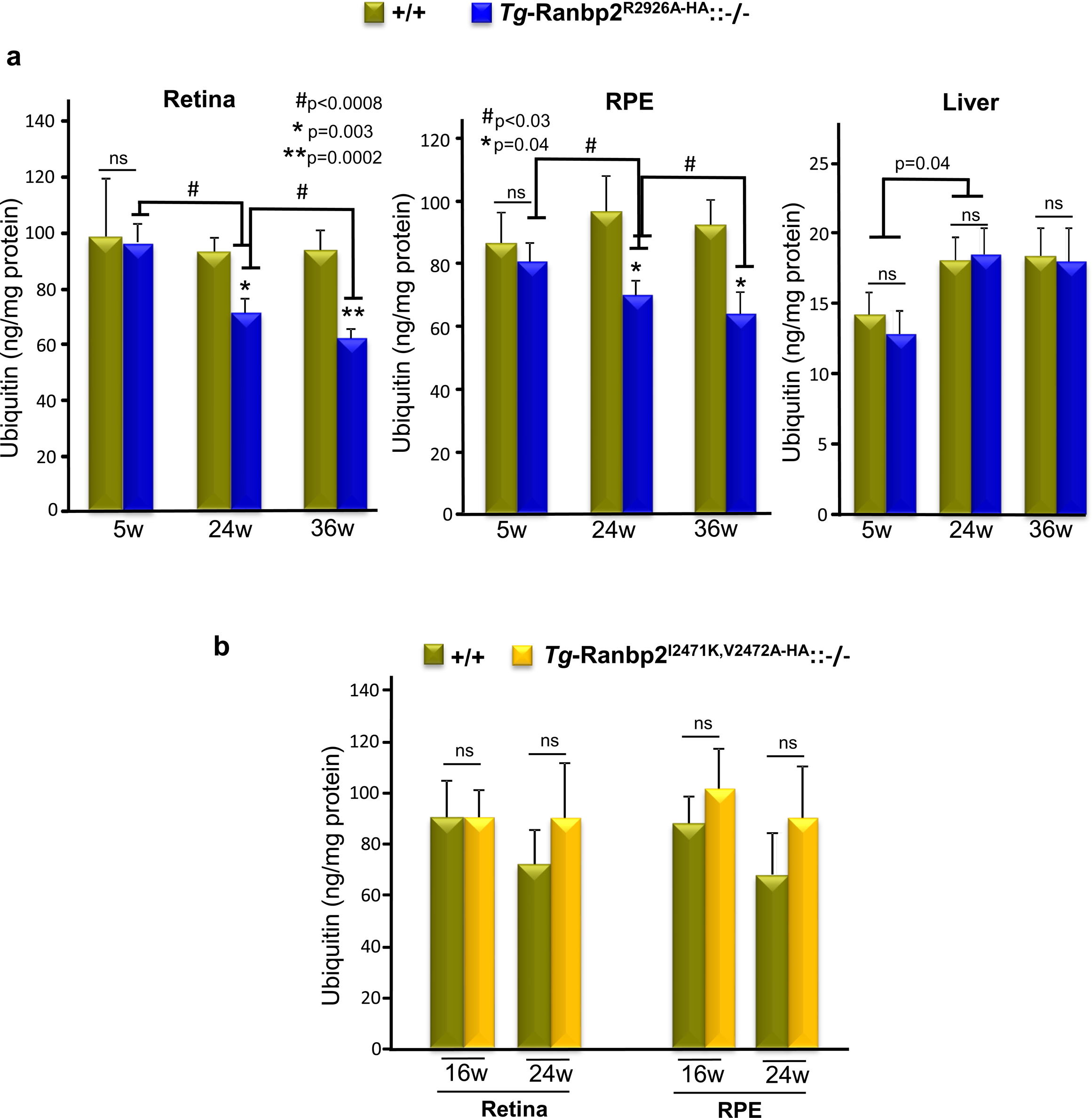
(a) Quantitation of total ubiquitin levels between retinae (left), RPE (middle) and liver (right) of wild type (+/+) and *Tg*-*Ranbp2^R2926A-HA^::-/-* mice at 5, 24 and 36-weeks of age. Total levels of ubiquitin significantly decline in retinae and RPE of *Tg*-*Ranbp2^R2926A-HA^::-/-* with aging. No changes of the total levels of ubiquitin in the liver are observed between genotypes with aging. There is a significant increase of the total level of ubiquitin in the liver between 5 and 24-week old mice of either genotype. Data represent the mean ± S.D (error bars), *n*=4 (5 and 24-week old mice), n=5 (+/+, 36-week old), n=3 (*Tg*-*Ranbp2^R2926A-HA^::-/-*, 36-week old). (b) Quantitation of total ubiquitin levels between retinae and RPE of wild type (+/+) and *Tg-Ranbp2^I2471K,V2472A-HA^::-/-* mice at 16 and 24-weeks of age. No significant differences are observed between genotypes at any age. Data represent the mean ± S.D (error bars), *n*=3. *Abbreviations*: w, weeks of age.

### Tg-Ranbp2^R2926A-HA^::-/- mice lack overt morphological changes of the chorioretina

We have previously shown that compared to wild type mice, 4-6 week old *Tg-Ranbp2^R2926A-HA^::-/-* mice neither develop overt morphological phenotypes of the chorioretina, nor changes in electrophysiological responses of outer retinal neurons (e.g. cone and rod photoreceptor neurons) [104]. In contrast, *Tg-Ranbp2^R2926A-HA^::-/-* display an improved response of the visual pathway compared to wild type mice that is reflected by a shorter delay of the implicit times of the dark-adapted visual evoked potentials (VEP) [104]. Hence, we examined the genotype-dependent effects of the decline of ubiquitylated substrates and changes in the proteostasis of neuroprotective substrates in the overall morphology of the chorioretina by light and ultrastructural microscopy of 9-month old wild type and *Tg-Ranbp2^R2926A-HA^::-/-* mice. As shown in figures 8a and 8b, there were no overt differences of the overall organization of chrorioretinal layers, RPE and retinal neurons between genotypes by light microscopy of retinal sections (figure 8a), nor by transmission electron microscopy (figure 8b). Hence, the proteostatic changes produced by the loss of PPIase and disturbance of C-terminal chaperone activity of CY of Ranbp2 do not affect the morphological organization of the chorioretina.

**Figure 8.**
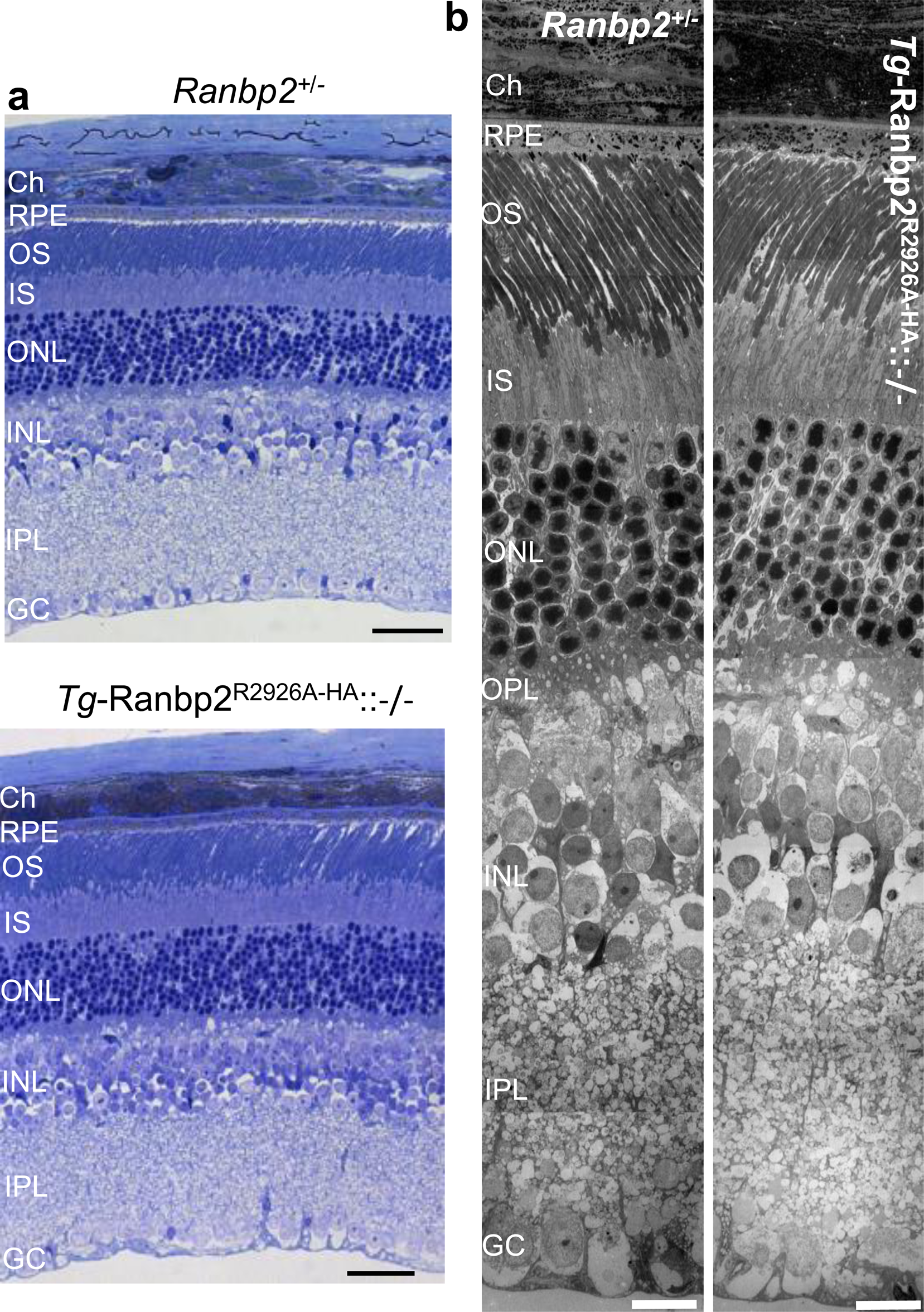
(a) Light photomicrographs of methylene blue-stained semithin sections of the central region of the chorioretina of *Ranbp2^+/-^*(top) and *Tg*-*Ranbp2^R2926A-HA^::-/-* mice (bottom) at 9-weeks of age. No overt morphological differences of the RPE and the retinal neurons are observed between the genotypes. Scale bar: 40 μm. (b) Ultrastructural photomicrographs of the central region chorioretina of *Ranbp2^+/-^* (left) and *Tg*-*Ranbp2^R2926A-HA^::-/-* mice (right) at 9-weeks of age. No overt differences are observed in the ultrastructure of the RPE and retinal neurons between the genotypes. Scale bar: 10 μm. *Abbreviations*: RPE, retinal pigment epithelium; OS, outer segments of rod photoreceptors; IS, inner segments of photoreceptors; ONL, outer nuclear layer (nuclei of photoreceptors), OPL, outer plexiform layer; INL, inner nuclear layer; IPL, inner plexiform layer; GC, ganglion cell layer; ch, choroid.

## DISCUSSION

This study identifies a critical role of the moonlighting activity of CY of Ranbp2 in the remodeling of the proteostasis of αA-crystallins and γ-crystallins. The expressions of these crystallins become down-regulated in post-weaning and naïve mice. Loss of PPIase and disturbance of C-terminal chaperone activities of CY of Ranbp2 selectively cause the respective up-regulation of αA-crystallin and the down-regulation of a subset of γ-crystallins in the retina and RPE. By contrast, a disturbance of the C-terminal chaperone activity alone of CY of Ranbp2 causes the down-regulation of a subset of γ-crystallins, but without affecting αA-crystallin proteostasis. Hence, loss of PPIase of CY of Ranbp2 alone underpins the increase of αA-crystallin. This opposite effect of CY of Ranbp2 on αA-crystallins and γ-crystallins is surprising, because γ-crystallins are reported to be substrates of the chaperone activity of αA-crystallin *in vitro* [34, 46–48] and likely *in vivo* [49–51]. Hence, the findings here presented uncover an independent functional relationship between αA-crystallin and γ-crystallins’ proteostasis by revealing that the proteostasis of αA-crystallins can be uncoupled from that of γ-crystallins by the loss of PPIase activity of CY of Ranbp2. These observations also raise the following implications.

First, γ-crystallins become the second known substrate of the C-terminal chaperone activity of CY of Ranbp2 besides L/M-opsin (figure 9a). The high molecular weight isoform - γ-crystallin^H^ may represent several γ-crystallin isoforms, as indicated by MS/MS of 2D-DIGE spots (figures 2, 4 and 5), because *i*) γ-crystallins comprise isoforms with a very close primary sequence homologies and molecular masses [115], and *ii*) the γ-crystallin antibody used is likely to cross-react with several isoforms. However, we cannot exclude that the C-terminal chaperone activity of CY of Ranbp2 has preferential activity towards one or more posttranslational modified isoforms of γ-crystallins and that are represented by γ-crystallin^H^. Regardless, these possibilities are not mutually exclusive. Future studies are needed to identify the exact isoforms and potential posttranslational modification(s) represented by γ-crystallin^H^. Second, the C-terminal chaperone activity of CY on γ-crystallin^H^ underscores the existence of another domain in CY of Ranbp2 on the opposite site of the PPIase pocket (figure 9a). A steric hindrance caused by the insertion of the 9-residue extension of the HA-tag at the C-terminal end of CY of Ranbp2 is likely to impair the chaperone activity of CY on γ-crystallins (figure 9a). These findings are consistent with genetic findings of missense and pathological mutations away from the PPIase pocket in selective cyclophilins and that affect their chaperone activity on restricted substrates [27, 29, 30, 118–120]. In this regard, our prior studies have also shown the existence of yet another domain in CY that associates with the neuroprotective substrate, STAT3, independently of the loss of PPIase and disturbance of C-terminal chaperone activities of CY of Ranbp2 [104]. CypB likely shares a similar domain with CY, because it also associates to STAT3 [121]. Third, the increase of αA-crystallin by loss of PPIase of CY of Ranbp2 supports that the PPIase activity acts as a prolyl *cis-trans* molecular switch on αA-crystallins that promotes their stability (e.g., oligomerization) or on a nucleocytoplasmic shuttling factor(s), which controls αA-crystallin expression. A role for the catalysis of peptidyl-prolyl *cis-trans* isomerization as a molecular and conformational switch in protein function has been established and proposed for the PPIase activity of CY of Ranbp2 and other cyclophilins as well for other PPIases [122–127]. In this regard, hnRNPA2B1 and HDAC4 are the only known two nucleocytoplasmic shuttling factor(s) that are regulated by the PPIase of CY; thus, they emerge as candidates to undergo conformational switches in the regulation of expression of αA-crystallins in non-lenticular tissues, such as the retina and RPE. Collectively with our prior studies, the findings support that the PPIase activity of Ranbp2 harnesses the activity of three substrates implicated in neuroprotection - αA-crystallin, HDAC4 and hnRNPA2B1 (figure 9A). Finally, a tug-of-war between the PPIase and chaperone (holdase) activities has been suggested for the PPIase pocket of cyclophilin A (CypA) in promoting and preventing the misfolding (aggregation) of α-synuclein by competition of the PPIase pocket of CypA with distinct sequences of α-synuclein [128]. Our results do not support a competition of αA-crystallin and γ-crystallin for the PPIase pocket of CY of Ranbp2. Instead, an analogous tug-of-war mechanism may involve the respective PPIase and-C-terminal chaperone activities of CY of Ranbp2 towards the regulation of αA-crystallin and γ-crystallin proteostasis by mass action shifts of these substrates.

**Figure 9.**
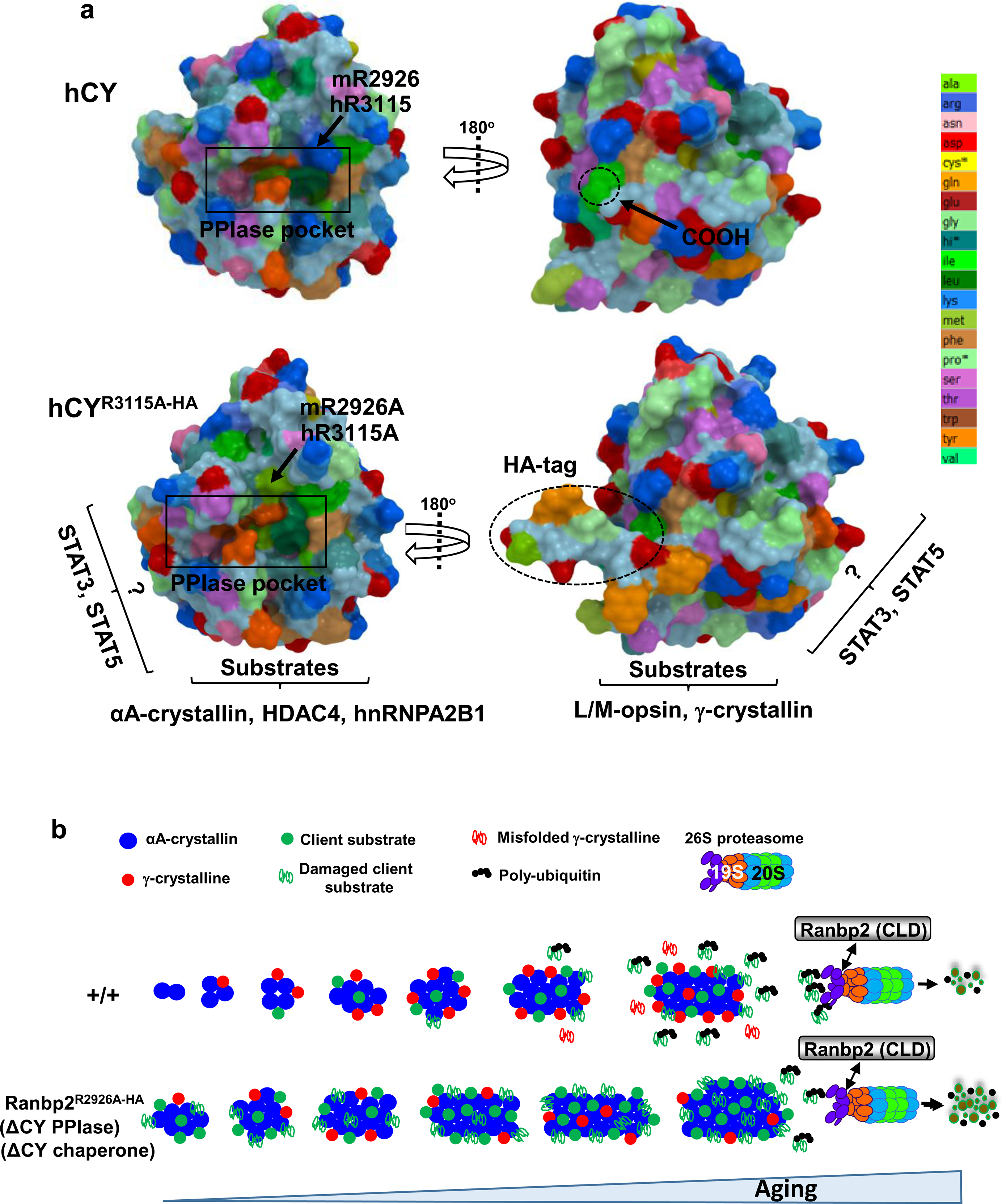
(a) Comparisons of molecular and geometric **s**urfaces of native and mutant isoforms of CY of Ranbp2 and substrates of CY. The top structures are representations of CY of human Ranbp2 (PDB: 4I9Y) depicting the side of the PPIase pocket (rectangular box) with the critical catalytic residue R3115 (or equivalent R2926 in *Mus musculus*; left) and on the opposite side, the surface harboring chaperone activity and the C-terminal end (right). The lower structures are representations of the mutant human isoform, CY^R3115-HA^ (equivalent to CY^R2926-HA^ in *Mus musculus*) with loss of PPIase catalytic activity caused by R3115A (R2926A in *Mus musculus*) (left) and with the insertion of an HA-tag at the C-terminal end of CY (right). The HA-tag causes the protrusion of the C-terminal end of CY and it affects the chaperone activity of this region likely by steric hindrance. The substrates affected by loss of PPIase activity and by the HA-tag disturbance of the C-terminal chaperone activity of CY are shown below the mutant structures. The regions of CY that chaperone latent and activated STAT3 and STAT5 do not overlap with the PPIase and C-terminal chaperone regions and remain molecularly undefined (shown with “?”). Surfaces of CY are color-coded by amino acid residues (right, colored strip). Mutant structure of CY was predicted with ColabFold against human CY and other cyclophilin templates. (b) Model of stimulation of proteostasis by loss of PPIase of CY of Ranbp2. Critical properties associated to αA-crystallin relate to its ability to convert to heterogeneous oligomers, to chaperone γ-crystallins and to act as holdases for misfolded/damaged client substrates. The chaperone capacity of αA-crystallin is limited and this capacity is compromised by chronic stress insults, such as aging, that promote the genesis of protein aggregates and ubiquitylated substrates that are resistant to UPS degradation or that overload the degraded capacity of the UPS with aging. Ranbp2 via its CLD (figure 1a) mediates the physical coupling to the 19S cap of the 26S proteasome and promotes substrate channeling. Loss of PPIase and disturbance of C-terminal chaperone activities of CY of Ranbp2 promote an increase of the chaperone capacity of αA-crystallin toward misfolded/damaged client substrates by an increase of αA-crystallin and a decrease of γ-crystallins, which are also susceptible to oxidative damage. This mass action shifts in αA-crystallin and γ-crystallins enhance the sequestration and promote the refolding of misfolded/damaged substrates by αA-crystallin and a decrease of the genesis of protein aggregates and ubiquitylated substrates, while preserving UPS function. These effects ultimately protect mice against molecular aging (e.g., proteotoxicity) and together with other neuroprotective substrates of CY (figure 9a), they preordain the chorioretina to neuroprotection against photo-oxidative stress.

The observation of the augmentation of αA-crystallin and γ-crystallins proteostasis also in mice harboring Ranbp2 with the mutations (I2471K, V2472A) in the SIM of Ranbp2 is surprising (figure 1a). These mutations overlap with CLD and IR1 of Ranbp2 (figure 1a) [110, 129] and suppress the binding of at least a SUMOylated substrate of Ranbp2, such as SUMOylated-RanGAP [130]. In the context of CLD function alone, the ectopic expression of CLD upregulates the proteostasis of ectopically expressed proteins, such Rpgrip1, whereas the I2471K and V2472A mutations abolish this effect. Another, key feature of CLD of Ranbp2 is that it specifically associates with a subcomplex of the 19S cap of the proteasome, likely via the S1/p112 (aka, Pmsd1) subunit, but the SIM mutations in CLD alone do not interfere with the binding of the 19S cap subunits to CLD [109, 110]. Further, the association of the 19S cap to CLD likely requires tissue-selective factors, because the binding activity is observed in the retina, but not in non-neuronal tissues [108, 109]. Physiologically, the SIM mutations in Ranbp2 appear to have a restricted substrate effect in the post-transcriptional decline of SUMOylated-RanGAP and HDAC4 proteostasis in the retina, and without affecting the UPS, such as the accumulation of ubiquitylated substrates, independently of the age of mice (figure 7b) [104].

Hence, prior and current findings support of a crosstalk between the PPIase-dependent activity of CY and CLD of Ranbp2 in regulation of proteostasis. Since αA-crystallin and HDAC4 are both physiological affected by perturbations of activities of CY and CLD of Ranbp2, αA-crystallin may serve as a tissue-selective factor, which modulates and bridges the presentation of client substrates by CY of Ranbp2 to the 19S cap of the 26S proteasome that associates to CLD of Ranbp2. HDAC4 may also regulate αA-crystallin expression or function by post-translational modification (e.g., deacetylation). Regardless, our findings add to emerging studies indicating that small HSPs and ATP-dependent HSPs play essential but complementary roles in bridging the disaggregation of misfolded proteins and their presentation to the UPS [131].

As noted earlier, insufficiency of Ranbp2 promotes the neuroprotection of the chorioretina against photo-oxidative stress with aging [11, 12]. Our prior studies and this work support that a deficit of CY PPIase activity of Ranbp2 is likely to contribute to the neuroprotection of the chorioretina against noxious stressors. To this effect, mice with loss of CY PPIase activity of Ranbp2 recapitulate molecular traits of haploinsufficiency of *Ranbp2*, such as a decline of ubiquitylated substrates against proteostatic stressors with aging. In line with these observations, a model emerges on yet another factor, αA-crystallin, which is regulated by the CY PPIase Ranbp2, and that is likely to preordain the chorioretina to neuroprotection against proteostatic stressors (figure 9b). In this model, up-regulation of αA-crystallin by loss of CY PPIase of Ranbp2 augments the proteostatic buffer capacity of the RPE and retina against misfolded or damaged substrates that accumulate with aging by enabling these substrates to refold or to be degraded by the UPS. Regardless, the ultimate outcome of this CY PPIase-dependent effect(s) is a decline of ubiquitylated-aggregates resistant to UPS degradation (figure 9b). The increase of the chaperone activity of αA-crystallin is attained by a mass action shift of αA-crystallin, whose increase promotes the dynamic oligomerization and poly-dispersion of αA-crystallin with an enhanced capacity to sequester misfolded or damaged substrates (figure 9b). This chaperone role is harnessed by the surveillance of misfolded/damaged proteins of the CY PPIase activity of Ranbp2, which with the physical coupling of its CLD to the 26S proteasome (figure 1a), together play critical roles in substrate channeling of ubiquitylated substrates and protein degradation.

The “foldase” activity linked to PPIases has been typically associated with the catalysis of a critical rate-limiting step of protein folding in the folding pathway of native proteins [132]. However, and in contrast to other studies [133, 134], emerging evidence, including some of our own findings, counter-intuitively indicate that PPIase activity promotes protein misfolding or aggregation of client substrates under selective environments possibly by lowering the energy barrier for protein misfolding [11, 12, 128] or by acting as a conformational switch in protein function [122–127]. Hence, the inhibition of PPIase activity toward selective client substrates emerges as a therapeutic tool in suppressing protein misfolding and aggregation. Negative allosteric sites affecting the PPIase activity of Ranbp2 CY and other cyclophilins [104, 135], small molecule inhibitors of CY PPIase of Ranbp2 [105] or post-translational modifications of the PPIase active site [136] emerge as novel therapeutic routes in suppressing protein misfolding and aggregation of client substrates, which underpin the pathogenesis of a spectrum of neurodegenerative diseases affecting the chorioretina and other neuronal tissues.

## MATERIALS & METHODS

### Generation of genetically engineered mice of Ranbp2

The production of genetically engineered lines of *Ranbp2* comprising RPE-cre::*Ranbp2^-/-^* [10], *Ranbp2^+/-^* [111] and transgenic mice (Tg) harboring bacterial artificial chromosomes (BAC) of the complete *Ranbp2* gene genetically engineered by BAC recombineering with a C-terminal HA-tag alone (*Tg*-Ranbp2^-HA^) [104] and with the mutations, W1889R and W2186R (*Tg*-Ranbp2^RBD2/3*-HA^) [10], I2471K and V2472A (*Tg*-Ranbp2^I2471K,V2472A-HA^) [104] or R2926A (*Tg*-Ranbp2^R2926A-HA^) [104] have been described. Hemizygous *Tg* alleles were placed and examined in RPE-cre::*Ranbp2^-/-^* or *Ranbp2^-/-^* genetic backgrounds. Experimental *Tg* lines of either sex were in a mixed C57Bl6/J background free of *rd1* and *rd8* alleles. Mice were reared in a pathogen-free transgenic barrier facility under 12-h light-dark cycles (6:00 AM - 6:00 PM), <70 lux, humidity and temperature-controlled conditions and with *ad libitum* access to water and chow diet 5LJ5 (Purina, Saint Louis, MO) at Duke University.

### Animal Study Protocols

Animal protocols were conducted as approved by the Institutional Animal Care and Use Committees of Duke University (A003-14-01). All procedures were in adherence to NIH guidelines for the care and use of laboratory animals, the National Academy of Sciences and the ARVO guidelines for the Use of Animals in Vision Research.

### Antibodies

Rabbit anti-hsc70 (1:3,000, ENZO Life Science, Farmingdale, NY, USA; Cat: ADI-SPA-816)), mouse anti-mHsp70 (1:600; Thermo Scientific, Rockford, IL; Cat: MA3–028), rabbit anti-COUP-TF1 [1:1,000 (IB), Abcam, Cambridge, MA, USA], rabbit anti-Nr2e3 (1:1,000, Proteintech; Cat: 14246-1-AP); rabbit anti-γ-crystallin and anti-βB2-crystallin (gift from Vasanth Rao, originated by Dr. Sam Zigler, NIH/NEI, Bethesda, MD); rabbit anti-αA-crystallin (gift from Vasanth Rao, originated by Joseph Horwitz, Jules Stein Eye Institute, UCLA); rabbit anti-GAPDH [1:500 (IB), Santa Cruz Biotechnology, Inc., Santa Cruz, CA, USA; Cat: sc-25778]; goat horseradish peroxidase-conjugated anti-rabbit IgG [25 ng/ml] were from Jackson ImmunoResearch Laboratories, Inc. (West Grove, PA).

### 2D-DIGE proteomics

RPE and retinal homogenates were solubilized in radio-immunoprecipitation assay buffer (RIPA) followed by buffer exchange in two-dimensional lysis buffer (7 M urea, 2 M thiourea, 4% CHAPS, 30 mM Tris-HCl, pH 8.8). 2-DIGE of RPE and retinal homogenates were labeled with CyDyes and resolved in analytical and Prep gels by Applied Biomics (Hayward, CA). Spots with differential expression between two or more genotypes with a twofold cut-off were identified and ranked with DeCyder “in-gel” analysis software. Spots of interest were isolated, trypsinized and the protein compositions of the spots were identified by mass spectrometry (MALDI-TOF/TOF) of mass spectra of tryptic peptides and by peptide fragmentation mapping. Combined MS and MS/MS spectra were used to search NCBI and SwissProt databases with the MASCOT search engine. Data analyses of samples were performed by the Ferreira laboratory and protein species with protein and total ion scores with confidence intervals (C.I.) of 100% were selected for independent experimental validation and analyses.

### Protein homogenates and immunoblot analyses

Tissue homogenate and immunoblot procedures were carried out as described elsewhere [10, 104, 108, 137]. Briefly, retinae and RPE were homogenized in radioimmune precipitation assay (RIPA) buffer with zirconium oxide beads (Next Advance, Averill Park, NY, USA, ZROB05) and a Bullet blender (Next Advance, BBX24) at 8000 rpm for 3 min. Protein concentrations of tissue homogenates were measured by the BCA method using BSA as the standard (Pierce™ BCA Protein Assay; Thermo Scientific, Cat: 23225). Equal amounts of homogenates (25 μg) were loaded in parallel with with pre-stained molecular weight markers and resolved by SDS-PAGE in premade 4–15% gradient Criterion gels (BioRad, Hercules, CA, USA). Resolved tissue homogenates were transferred onto polyvinylidene difluoride (PVDF), blocked with non-fat dry milk block solution (Bio-Rad, Hercules, CA), incubated with primary antibodies and goat horseradish peroxidase-conjugated anti-rabbit IgG. Blots were reprobed for Hsc70, mHsp70 or Gapdh for normalization and quantification of proteins. Immunoreactive proteins were visualized with enhanced chemiluminescence reagent (ThermoScientific, Cat: 32106) followed by exposure to X-ray Hyperfilm (Amersham Biosciences). Unsaturated band intensities were quantified by densitometry with Metamorph v7.0 (Molecular Devices, San Jose, CA, USA), and integrated density values (idv) of bands were normalized to the background and idv of loading/housekeeping controls.

### Measurement of ubiquitylated proteins

The UbiQuant ELISA kit as directed by the manufacturer (LifeSensors, Malvern, PA) was used to determine the absolute and total amount of ubiquitin and ubiquitylated proteins in the RPE, retinae and liver between genotype and ages.

### Semithin sections and light and transmission electron microscopy

The preparation of specimens for light and transmission electron microscopy (TEM) microscopy were as described in prior studies [12, 13, 116, 137, 138]. Briefly, mice were anesthetized and perfused, and eyes were fixed with 2.0% glutaraldehyde and 2%paraformaldehyde (1:1) in 0.1 M sodium cacodylate buffer, pH 7.4 overnight at 4 °C. For light microscopy, semi-thin sections (0.5 μm) of posterior eyecups along the vertical meridian were mounted and stained with 1% methylene blue. Images were captured with an Axiopan-2 light microscope coupled to an AxioCam HRc digital camera (Carl Zeiss, Germany). For ultrastructural microscopy by TEM, specimens were post-fixed in 2% osmium tetraoxide in 0.1% cacodylate buffer, embedded in Spurr resin and ultrathin sections (60 nm-thick) were collected with a Leica Ultracut S (Leica Microsystems, Waltzer, Germany). Sections were stained with 2% uranyl acetate and 4% lead citrate and examined on a Phillips BioTwin CM120 electron microscope equipped with Gatan Orius and Olympus Morada digital cameras.

### Molecular modeling of mutant CY structure of human Ranbp2

The mutation, R3115A (equivalent to R2926A in *Mus musculus*) with and without a C-terminal human influenza hemagglutinin (HA)-tag in CY were introduced to the primary sequence of Ranbp2 CY and whose structure was determined at 1.75 Å (PBD 4I9Y) [139]. The HA-tag has been shown not to assume a well-defined secondary structure [140]. Then, the predicted three-dimensional (3D) structure of mutant CY was determined against orthologue cyclophilin templates in PBD, including human CY of Ranbp2 (PBD: 4I9Y), by using AlphaFold2 and structure energy minimization (Open MM) with ColabFold [141]. When applicable, predicted structures were imported to ICM software (MolSoft LLC, San Diego) for publication rendering [142].

### Statistical analyses

Samples sizes/independent biological replicates were collected (power > 0.8) and these were comparable with other studies using the same mouse lines and genotypes. Student’s t-test for two groups was used. Data are reported as average values ± SD. Differences between groups were considered statistically significant when p-value < 0.05.

### CRediT authorship contribution statement

Hemangi Patil: Investigation (lead); Formal Analysis; Validation; Kyoungin Cho: Investigation (supporting); Formal Analysis; Validation; Paulo A Ferreira: Conceptualization; Methodology; Writing - original draft, Writing - review & editing; Funding Acquisition; Project Administration.

## ACKNOWLEDGEMENTS.

This work was supported by the National Institutes of Health Grants EY019492 and GM083165 to PAF.

